# *De novo* variants in population constrained fetal brain enhancers and intellectual disability

**DOI:** 10.1101/621029

**Authors:** Matias G De Vas, Myles G Garstang, Shweta S Joshi, Tahir N Khan, Goutham Atla, David Parry, David Moore, Ines Cebola, Shuchen Zhang, Wei Cui, Anne K Lampe, Wayne W Lam, David R FitzPatrick, Jorge Ferrer, Madapura M Pradeepa, Santosh S Atanur

## Abstract

**Purpose:** The genetic aetiology of a major fraction of patients with intellectual disability (ID) remains unknown. *De novo* mutations (DNMs) in protein-coding genes explain up to 40% of cases, but the potential role of regulatory DNMs is still poorly understood.

**Methods:** We sequenced 70 whole genomes from 24 ID probands and their unaffected parents and analyzed 30 previously sequenced genomes from exome-negative ID probands.

**Results:** We found that DNVs were selectively enriched in fetal brain-specific enhancers that show purifying selection in human population. DNV containing enhancers were associated with genes that show preferential expression in the pre-frontal cortex, have been previously implicated in ID or related disorders, and exhibit intolerance to loss of function variants. DNVs from ID probands preferentially disrupted putative binding sites of neuronal transcription factors, as compared to DNVs from healthy individuals and most showed allele-specific enhancer activity. In addition, we identified recurrently mutated enhancer clusters that regulate genes involved in nervous system development (*CSMD1*, *OLFM1* and *POU3F3)*. Moreover, CRISPR-based perturbation of a DNV-containing enhancer caused *CSMD1* overexpression and abnormal expression of neurodevelopmental regulators.

**Conclusion:** Our results, therefore, provide new evidence to indicate that DNVs in constrained fetal brain-specific enhancers play a role in the etiology of ID.

## Introduction

Intellectual disability (ID) is a neurodevelopmental disorder that is characterized by limitations in both intellectual functioning and adaptive behavior^1^. The clinical presentation of ID is heterogeneous, often coexisting with congenital malformations or other neurodevelopmental disorders such as epilepsy and autism^1^, and the worldwide prevalence is thought to be near 1%^2^. Throughout the past decade, great progress has been made in identifying the genetic causes of ID. *De novo* protein truncating variants and copy number variants affecting a large set of protein-coding genes explain up to 40% of ID cases^1, 3, 4^, whereas the genetic etiology of a major fraction of ID patients still remains unknown. *De novo* variants (DNV) in non protein-coding regions of the genome could plausibly explain some of the cases in which no causal coding variant has been identified.

Previous studies have implicated variants in long-range transcriptional regulatory elements, also known as enhancers, in monogenic developmental disorders including preaxial polydactyly (*SHH*)^5, 6^, Pierre Robin sequence (*SOX9*)^7^, congenital heart disease (*TBX5*)^8^ and pancreatic agenesis (*PTF1A*)^9^. A systematic analysis of variants in evolutionarily ultra-conserved regions of the genome has estimated that around 1-3% of the developmental disorder patients without disease-causing coding variants would carry disease-causing DNVs in fetal brain-active regulatory elements^10^. A large-scale genome sequencing of patients with autism spectrum disorder (ASD) demonstrate that DNVs in conserved promoter regions contribute to ASD, while no significant association was found between enhancer variants and ASD^11^.

Despite these precedents, the efforts to implicate enhancer variants in human disease, however, face numerous challenges. Importantly, it is currently not possible to readily discern functional enhancer variants from non-functional or neutral variants based on sequence features. This can be partially addressed through experimental analysis of regulatory DNA variants. Moreover, we still need a full understanding of which regulatory regions and which subsequences within the regulatory regions are most likely to harbor disruptive variants. In addition, one of the biggest challenges in interpreting variants in regulatory regions is to correctly associate regulatory regions to the potential target genes. Systematic identification of tissue specific promoter-enhancer interaction maps would help identification of regulatory regions that are associated with disease relevant genes.

To address some of these challenges, we have sequenced genome of patients with ID, and examined whether DNVs target a subset of regulatory regions that are most likely to harbor etiological defects. We hypothesized that ID could result from DNVs in enhancers that are specifically active during brain development. We further reasoned that although evolutionary conservation is an important metric to prioritize genomic regions, advanced human cognition has been attributed to human fetal brain enhancers that are gained during evolutionary expansion and elaboration of the human cerebral cortex^12^, hence critical regulatory sequences for intellectual functions may show sequence constraints within human populations regardless of their evolutionary conservation.

In this study, we deployed whole genome sequence analysis, integrative genomic, epigenomic and experimental studies to show that DNVs in patients with ID are selectively enriched in fetal brain-specific enhancers and human brain gained enhancers that exhibit sequence constraint within human populations. We further show that such DNVs map to enhancers that are associated with known ID genes, genes that are intolerant to variants and genes specifically expressed in the pre-frontal cortex. Furthermore, we identify three fetal brain-specific enhancers domains with recurrent DNVs, and provide experimental evidence that candidate variants alter enhancer activity in neuronal cells. These results provide new level of evidence that supports a role of DNVs in neurodevelopmental enhancers in the aetiology of ID.

## Materials and methods

### Selection criteria of intellectual disability patients for this study and ethical approval

The inclusion criteria for this study were that the affected individuals had a severe undiagnosed developmental or early onset pediatric neurological disorder and that samples were available from both unaffected parents. Written consent was obtained from each patient family using a UK multicenter research ethics approved research protocol (Scottish MREC 05/MRE00/74).

### Sequencing and quality control

Genome sequencing was performed on the Illumina X10 at Edinburgh Genomics. Genomic DNA (gDNA) samples were evaluated for quantity and quality using an AATI, Fragment Analyzer and the DNF-487 Standard Sensitivity Genomic DNA Analysis Kit. Next Generation sequencing libraries were prepared using Illumina SeqLab specific TruSeq Nano High Throughput library preparation kits in conjunction with the Hamilton MicroLab STAR and Clarity LIMS X Edition. The gDNA samples were normalized to the concentration and volume required for the Illumina TruSeq Nano library preparation kits then sheared to a 450bp mean insert size using a Covaris LE220 focused-ultrasonicator. The inserts were ligated with blunt ended, A-tailed, size selected, TruSeq adapters and enriched using 8 cycles of PCR amplification. The libraries were evaluated for mean peak size and quantity using the Caliper GX Touch with a HT DNA 1k/12K/HI SENS LabChip and HT DNA HI SENS Reagent Kit. The libraries were normalised to 5nM using the GX data and the actual concentration was established using a Roche LightCycler 480 and a Kapa Illumina Library Quantification kit and Standards. The libraries were normalised, denatured, and pooled in eights for clustering and sequencing using a Hamilton MicroLab STAR with Genologics Clarity LIMS X Edition. Libraries were clustered onto HiSeqX Flow cell v2.5 on cBot2s and the clustered flow cell was transferred to a HiSeqX for sequencing using a HiSeqX Ten Reagent kit v2.5.

### Alignment and variant calling

The de-multiplexing was performed using bcl2fastq (2.17.1.14) allowing 1 mismatch when assigning reads to a barcodes. Adapters were trimmed during the de-multiplexing process. Raw reads were aligned to the human reference genome (build GRCh38) using the Burrows-Wheeler Aligner (BWA) mem (0.7.13)^13^. The duplicated fragments were marked using samblaster (0.1.22)^14^. The local indel realignment and base quality recalibration was performed using Genome Analysis Toolkit (GATK; 3.4-0-g7e26428)^15–17^. For each genome SNVs and indels were identified using GATK (3.4-0-g7e26428) HaplotypeCaller^18^ creating a gvcf file for each genome. The gvcf files of all the individuals from the same family were merged together and re-genotyped using GATK GenotypeGVCFs producing single VCF file per family.

### Variant filtering

Variant Quality Score Recalibration pipeline from GATK^15–17^ was used to filter out sequencing and data processing artifacts (potentially false positive SNV calls) from true SNV and indel calls. First step was to create a Gaussian mixture model by looking at the distribution of annotation values of each input variant call set that match with the HapMap 3 sites and Omni 2.5M SNP chip array polymorphic sites, using GATK VariantRecalibrator. Then, VariantRecalibrator applies this adaptive error model to both known and novel variants discovered in the call set of interest to evaluate the probability that each call is real. Next, variants were filtered using GATK ApplyRecalibration such that final variant call set contains all the variants with 0.99 or higher probability to be a true variant call.

### De novo variants (DNV) calling and filtering

The de novo variants (DNVs) were called using GATK Genotype Refinement workflow. First, genotype posteriors were calculated using sample pedigree information and the allele accounts from 1000 genome sequence data as a prior. Next, the posterior probabilities were calculated at each variant site for each sample of the trio. Genotypes with genotype quality (GQ) < 20 based on the posteriors are filtered out. All the sites at which both the parents genotype was homozygous reference (0/0) and child’s genotype was heterozygous (0/1), with GQs >= 20 for each sample of the trio, were annotated as the high confidence DNVs. Only high confident DNVs that were novel or had minor allele frequency less than 0.0001 in 1000 genomes project were selected for further analysis.

Because, the majority of the publically available datasets including epigenomic datasets are mapped to human genome assembly version hg19, we lifted over all the DNV co-ordinates to hg19 using liftover package. All the variant co-ordinates presented in this paper are from hg19 human genome assembly.

### DNV annotations

DNV annotations were performed using Annovar^19^. To access DNV location with respect to genes, refseq, ENSEMBL and USCS annotations were used. To determine allele frequencies, 1000 genome, dbSNP, Exac and GnomAD databases were used. To determine pathogenicity of coding DNVs, annotations were performed with CADD, DANN, EIGAN, FATHMM and GERP++ pathogenicity prediction scores. In addition, we determined whether any coding DNV has been reported in ClinVar database as a disease-causing variant.

### Structural variant detection and filtering

To detect structural variants (SV), we used four complimentary SV callers: BreakDancer v1.3.6 ^20^, Manta v1.5.0^21^, CNVnator v0.3.3^22^ and CNVkit v0.9.6^23^. The BreakDancer and Manta use discordant paired end and split reads to detect deletions, insertions, inversions and translocations, while CNVnator and CNVkit detect copy number variations (deletions and duplications) based on read-depth information. The consensus SV calls were derived using MetaSV v0.4^24^. The MetaSV is the integrative SV caller, which merges SV calls from multiple orthogonal SV callers to detect SV with high accuracy. We selected SVs that were called by at least two independent SV callers out of four.

To detect *de-novo* SV, we used SV2 v1.4.1^25^. SV2 is a machine-learning algorithm for genotyping deletions and duplications from paired-end genome sequencing data. In *de novo* mode SV2 uses trio information to detect *de novo* SVs at high accuracy.

### Tissue specific enhancer annotations

Roadmap Epigenomic Project^26^ chromHMM segmentations across 127 tissues and cell types were used to define brain-specific enhancers. We selected all genic (intronic) and intergenic enhancers (“6_EnhG and 7_Enh) from male (E081) and female fetal brain (E082). This was accomplished using genome-wide chromHMM chromatin state classification in rolling 200bp windows. All consecutive 200bp windows assigned as an enhancer in fetal brain were merged to obtain enhancer boundaries. A score was assigned to each enhancer based on the total number of 200bp windows covered by each enhancer. Next, for each fetal brain enhancer, we counted the number of 200bp segments assigned as an enhancer in the remaining 125 tissues and cell types. This provided enhancer scores across 127 tissues and cell types for all fetal brain enhancers. To identify fetal brain specific enhancers, Z scores were calculated for each fetal brain enhancer using the enhancer scores. Z scores were calculated independently for male and female fetal brain enhancers. Independent Z score cutoffs were used for both male and female fetal brain enhancers such that approximately 35% of enhancers were selected. To define open accessible chromatin regions within brain-specific enhancers, we intersected enhancers with DNAse-seq data from Roadmap Epigenomic Project^26^ from male (E081) and female fetal brain (E082) respectively. Next, the male and female fetal brain specific enhancers were merged together to get final set of 27,420 fetal brain specific enhancers. We used similar approach to identify tissue specific enhancers for selected fetal and adult non-brain tissues.

### Human gain enhancers

Human gain enhancers published previously by Reilly et al^12^ were downloaded from Gene Expression Omnibus (GEO) using accession number GSE63649.

### *De novo* variants from healthy individuals

We downloaded *de novo* variants identified in healthy individual in genomes of the Netherland (GoNL) study^27^ from GoNL website.

### Fetal brain-specific genes

Roadmap Epigenomic Project^26^ gene expression (RNA-seq) data from 57 tissues was used to identify fetal brain-specific genes. We used female fetal brain gene expression data, as RNA-seq data was available only for female fetal brain. For each gene, Z scores were calculated using RPKM values across 57 tissues. The genes with Z score greater than two were considered as the brain specific genes.

### *De novo* variant enrichment analysis

The expected number of *de novo* variants (DNVs) in fetal brain specific enhancers and human gain enhancers was estimated using the previously defined framework for *de novo* variants^28^. The framework for the null variant model is based on tri-nucleotide context where the second base is mutated. Using this framework, the probability of variant for each enhancer was estimated based on the DNA sequence of the enhancer. Probability of variant of all the enhancers within the enhancer set (fetal brain specific enhancers and human gain enhancers) was summed to estimate the probability of variant for the entire enhancer set. The probability of variant for fetal brain specific enhancers and human gain enhancers was estimated separately. To estimate the expected number of DNVs, the probability of variant for each enhancer set was multiplied by the cohort size (n=47). To estimate the significance of observed number of DNVs over expected number, Poisson distribution probabilities were invoked using R function ppois.

### Enrichment of recurrently mutated enhancer clusters

The enhancer clusters were randomly shuffled 1000 times. For each iteration we estimated number of enhancer clusters with more than one variant. Then we counted the number of times more than or equal to two variants were observed in three or more enhancer clusters. This number was then divided by 1000 to calculate P-value.

### DNV effect on transcription factor binding

The R bioconductor package motifbreakR^29^ was used to estimate the effect of DNV on transcription factor binding. The motifbreakR works with position probability matrices (PPM) for transcription factors (TF). MotifbreakR was run using three different TF databases: viz. homer, encodemotif and hocomoco. To avoid false TF binding site predictions, either with reference allele or with alternate allele, a stringent threshold of 0.95 was used for motif prediction. DNVs that create or disturb a strong base (position weight >=0.95) of the TF motif, as predicted by motifbreakR, were selected for further analysis.

### Prediction of target genes of enhancers

Three different methods were used to predict the potential target genes of enhancers.

Chromosome conformation capture (Hi-C) comprehensively detects chromatin interactions in the nucleus; however, it is challenging to identify individual promoter-enhancer interactions using Hi-C due to the complexity of the data. In contrast, promoter capture Hi-C (PCHi-C) specifically identifies promoter-enhancer interactions as it uses sequence capture to enrich the interactions involving promoters of annotated genes^30^. The significant interactions between promoters and enhancers identified using PCHi-C in neuronal progenitor cells^31^ were used to assign target genes to the DNV containing enhancers. The enhancers were overlapped with the PCHi-C HindIII fragments. If an overlap was found between an enhancer and the PCHi-C HindIII fragment, the significantly interacting regions (PCHi-C HindIII fragments representing promoters of the genes) of the PCHi-C HindIII fragment were extracted to assign genes to the enhancers.

For an enhancer to interact with a promoter, both promoter and enhancer need to be active in specific cells at a specific stage. To identify promoter-enhancer interactions, all the active promoters in fetal brain (as defined by chromHMM segmentation) were extracted. Promoter-enhancer interactions occur within topologically associated domains (TAD) hence, promoters that were located within the same TAD as that of a DNV containing enhancer were used for analysis.

For each enhancer and promoter, H3K27ac counts were extracted from all tissues for which H3K27ac data was available in the Roadmap Epigenomic Project^26^ ChIP-seq dataset. For fetal brain, H3K27ac ChIP-seq data published by Reilly *et al*^12^ was used because H3K27ac ChIP-seq data was not available in Roadmap Epigenomic Project ChIP-seq dataset for fetal brain. The Spearman rank correlation coefficient (Spearman’s rho) was calculated between each enhancer-promoter pair within the TAD using Scipy stats.spearmanr function from Python. The pervariant test was performed to identify the significance of the correlation. The counts were randomly shuffled, independently for enhancers and promoters, 1000 times to calculate an adjusted *P* value. The interactions with an adjusted *P* value less than 0.01 were considered as a significant interaction between the enhancer and promoter.

Finally, if any enhancer remained unassigned to a gene using these approaches, they were assigned to fetal brain expressed genes within the TAD. A gene with an expression level more than or equal to 1 TPM in the Roadmap Epigenomic Project fetal brain RNA-seq data was considered to be expressed in the fetal brain.

### Gene enrichment analysis

To test if enhancer associated genes were enriched for genes previously implicated in neurodevelopmental disorders, three different gene sets were used. 1) Intellectual disability (ID) gene list published in the review by Vissers *et al*^1^ was downloaded from Nature website 2) We compiled all the genes implicated in neurodevelopmental disorders in Deciphering Developmental Disorder (DDD) project^32^. 3) All the genes implicated in autism spectrum disorder were downloaded from SFARI browser. Significance of enrichment was tested using hypergeometric test in R.

Gene ontology enrichment and tissue enrichment analysis was performed using web-based tool Enricher (http://amp.pharm.mssm.edu/Enrichr/).

Probability of loss of function intolerance (pLI) scores for each gene was downloaded from Exome Aggregation Consortium (ExAC) browser (http://exac.broadinstitute.org/). Significance of enrichment was tested using hypergeometric test in R.

### Cell culture

LUHMES cells were purchased from ATCC (CRL-2927). Cells were cultured as previously described (Scholz et al, JNC 2011). Briefly, cells were attached on pre-coated multi-well plates (50 ug/mL poly-L-ornithine and 1 ug/mL fibronectin - Sigma) and grown in proliferation medium: Advanced DMEM/F-12 plus N-2 Supplement (Thermo Fisher), 2 mM L-glutamine (Sigma) and 40 ng/mL recombinant basic fibroblast growth factor (R&D Systems). Cells were differentiated into neurons for 7 days using standard differentiation medium: Advanced DMEM/F-12 plus N-2 Supplement, 2 mM L-glutamine, 1 mM dibutyryl cAMP (Sigma), 1 ug/mL tetracycline (Sigma) and 2 ng/mL recombinant human GDNF (R&D Systems).

Neuroblastoma cell line (SH-SY5Y) was maintained in DMEM/F12 media (Gibco), 1% penicillin-streptomycin, 10 % fetal bovine serum and 2 mM L-glutamine.

### Dual luciferase enhancer assays

Enhancer and control regions (500-600 bp) were amplified from human genomic DNA from HEK293T cells using Q5 High-Fidelity Polymerase (NEB). Amplified fragments were cloned into pGL4.23 plasmid (Promega), which consists of a minimal promoter and the firefly luciferase reporter gene. These regions were mutagenized in order to introduce the *de novo* variants of interest using the Q5 Site-Directed Mutagenesis kit (NEB) using non overlapping primers. pGL4.23 plasmids containing putative enhancer DNA were sequence verified and transfected, together with a Renilla luciferase expressing vector (pRL-TK Promega) into SHSY-5Y cells using Lipofectamine 3000 (Invitrogen) following manufacturer’s protocol. Firefly and Renilla luciferase activity was measured 24 hours after transfection using the Dual-Luciferase Reporter Assay System (Cat. number E1910, Promega) as per the manufacturer’s instructions. Primers used to amplify genomic DNA and for mutagenesis are provided in **Table S11**.

### CRISPR interference lentivirus preparation and transduction

To generate viral particles, 293FT cells were transfected with 3^rd^ generation lentiviral plasmids (pMDLg/pRRE, pRSV-Rev and pMD2.G; Addgene #12251,12253 and 12259, respectively) along with a plasmid encoding KRAB-dCAS9 (Addgene 118155) using the PEIpro reagent (Polyplus-Transfection). Culture medium was changed the day after transfection (day 1) and virus was collected on day 3. Lentiviral particles were concentrated using Lenti-X Concentrator (Takara). LUHMES cells were seeded in 12-well plates and immediately transduced with 30 µl of virus concentrate. Culture medium was changed the following day and selection (4.5 µg/mL blasticidin) was started 3 days after transduction and continued until control untransduced cells were completely dead. A list of oligonucloetides used for sgRNA cloning is provided in **Table S12.**

### Genomic DNA extraction

Genomic DNA was extracted by a modified version of the salting-out method. Briefly, cells were lysed in Lysis Buffer (100 mM Tris-HCl pH 8.5; 5 mM EDTA; 200 mM NaCl; 0.2 % SDS) plus 4 U/mL of Proteinase K (Thermo Fisher) for at least 2 hours at 55°C with agitation. Then, 0.4x volumes of 5 M NaCl were added to the mixture and centrifuged at max. speed for 10 min. DNA in the supernatant was precipitated with 1x volume of isopropanol. After centrifugation, the pellet was washed with 70% ethanol and air dried for half an hour. DNA was resuspended in water and incubated for at least one hour at 37°C with agitation.

### RNA isolation, cDNA synthesis and RT-qPCR

RNA was extracted using the RNeasy Mini Kit (QIAGEN) and cDNA was produced with the Transcriptor First Strand cDNA Synthesis Kit (Roche). RT-qPCR reactions were performed with SYBR Green Master Mix (Thermo Fisher) and run on an Applied Biosystems QuantStudio 12K Flex Real-Time PCR machine. Relative gene expression values were calculated with the –ΔCt method, using TATA-Box Binding Protein (TBP) as a house-keeping gene for normalization. Oligonucleotides used for qPCR are provided in **Table S13**.

### Statistical analysis

All luciferase experiments and gene quantification using qPCR were done in biological triplicates. The significance level was calculated using two-tailed t-test.

### RNA-seq and data analysis

RNA-seq libraries using RNA extracted from control LUHMES cells, LUHMES cells with CSMD1 enhancer CRISPRi, control differentiated neurons and CRISPRi differentiated neurons were generated in triplicate. RNA-seq libraries were sequenced on one lane of Illumina Hi-seq 4000 with 75bp paired end sequencing.

Sequencing data for RNA-Seq samples are adapter trimmed using Fastp and mapped against a human reference genome GRCh38 using splice aware aligner STAR v2.6.1^33^ with GENCODE v29 gene annotations (https://www.gencodegenes.org/human/release_29.html). We generated raw counts per gene using the FeatureCounts tool (v1.6.3)^34^. The differential expression analysis was performed using DESeq2 v1.24.0^35^.

## Results

### Genome sequencing and identification of *de novo* variants

We performed genome sequencing (GS) of 70 individuals including 24 probands with severe intellectual disability (ID) and their unaffected parents at an average genome-wide depth of 37X **(Table S1)**. Our cohort includes 22 trios and one quad family with two affected probands. We identified on average 4.08 million genomic variants per individual that include 3.36 million single nucleotide variants (SNVs) and 0.72 million short indels **(Table S1)**. We focused our analysis on *de novo* variants (DNVs), as it has been shown that DNVs contribute significantly to neurodevelopmental disorders^32, 36^. We identified a total of 1,261 DNVs in 21 trios after excluding one trio from the analysis due to an excessively high number of DNVs. An average of 60 high quality DNVs per proband were identified, which includes 55.2 SNVs and 4.8 indels per proband **(Table S2)**. The number of DNVs identified in this study is similar to the number of DNVs identified per proband in previous GS studies on neurodevelopmental disorders^4, 11, 37^. It has been shown that *de novo* copy number variants (CNVs) play a significant role in severe ID^4^. We identified a total of three *de novo* CNVs in our ID probands (**Table S3**).

### Protein-coding *de novo* variants and copy number variants

The role of protein truncating variants in ID is well established. Hence, we first looked at DNVs located in protein-coding regions of the genome. A total of 23 DNVs were located in protein-coding regions (average 1.1 DNVs per proband). Of the 23 coding variants, 15 were non-synonymous coding variants or protein truncating variants. In six ID probands, we identified various types of potentially disease-causing variants in the genes *KAT6A*, *TUBA1A*, *KIF1A*, *NRXN1* and *PNKP,* all of them previously implicated in ID^1, 4^. The variant in *KAT6A* resulted in a premature stop codon while genes *TUBA1A* and *KIF1A* showed non-synonymous coding variants, which have been reported as a likely disease-causing and disease-causing respectively in ClinVar^38^ **(Table S4)**. One *de novo* CNV resulted in partial deletion of *NRXN1,* a known ID gene. A family with two affected siblings was analyzed for the presence of recessive variants. We identified a homozygous 17bp insertion in the gene *PNKP* **(Table S4)** in both siblings. This insertion has been reported as disease-causing in ClinVar^38^. These findings confirmed the disease-causing role of DNVs in ID.

### ID *de novo* variants are preferentially located in constrained fetal brain enhancers

In our severe ID cohort, we did not identify disease-causing coding DNVs in 17 ID cases (∼70%), and hence decided to investigate potentially disease-causing variants in disease-relevant enhancer regions. We further analysed 30 previously published severe ID samples in which no disease-causing protein-coding DNVs have been found using GS^4^, yielding a total of 47 exome-negative ID cases.

We hypothesized that DNVs in fetal brain-specific enhancers could perturb expression levels of genes that are essential for brain development, leading to ID. We therefore identified 27,420 fetal brain-specific enhancers using the data from Roadmap Epigenomics project^26^ (see Methods). The majority (76.52%) of these fetal brain specific enhancers were found to be candidate cis-Regulatory Elements (ccREs) defined by ENCODE3^39^. In addition, we analyzed 8,996 human brain gained enhancers that have been shown to be active during cerebral corticogenesis^12^. All the downstream analysis was performed on a combined set of fetal brain-specific enhancers and human fetal brain gained enhancers, which we refer as **fetal brain enhancers** throughout the manuscript.

A total of 83 DNVs (an average of 1.77 DNVs per proband) were located within fetal brain enhancers. It has been shown that the DNVs from individuals with neurodevelopmental disorders have been enriched in fetal brain active conserved non-coding elements^10^. Hence, we investigated enrichment of observed number of DNVs over expected in our ID cohort in multiple fetal and adult tissue specific enhancers including fetal brain and adult brain subsections using previously defined framework for interpreting DNVs^28^. As a control we used a set of randomly selected, sequence composition matched quiescent regions and the DNVs identified in healthy individuals in the Genome of Netherlands (GoNL)^27^. After applying multiple test correction we did not find a significant enrichment for DNVs in any tissue specific enhancers tested (**Figure 1a)**. However, fetal brain enhancers showed the largest excess of DNVs as compared to the expected number of variants (**Figure 1a**). Other tissues showed expected or fewer DNVs compared to the expected while adult brain specific enhancers showed a depletion of DNVs (**Figure 1a**). This result was consistent with the expectation that variants in enhancers that are active during fetal brain development play a role in ID, which is a severe early-onset neurodevelopmental phenotype, rather than variants in enhancers that are active in adult brain or other tissues.

**Figure 1:**
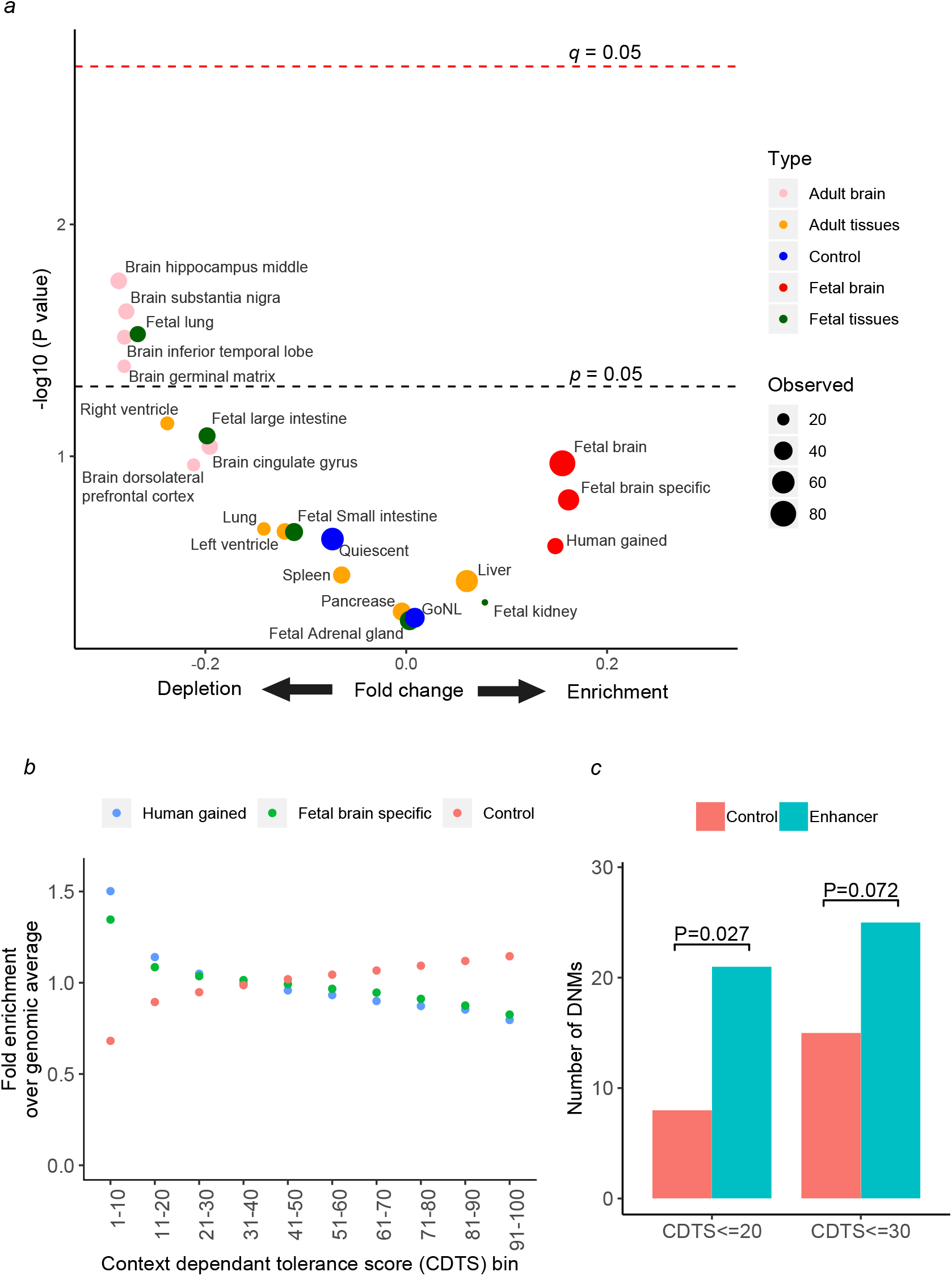
Enrichment of *de novo* variants (DNVs) in constrained enhancers. a) Enrichment or depletion of observed number of DNVs over expected number across multiple tissues and controls. The x-axis scale is centred around zero which indicates an equal number of observed, and expected number, of DNVs. Positive numbers of x-axis indicate enrichment and negative numbers indicate depletion of DNVs. The red dots indicate fetal brain enhancers while pink dots indicate enhancers from adult brain subsections. Green dots indicate tissue specific enhancers from fetal tissues other than brain while orange dots indicate tissue specific enhancers from adult tissues. Two controls; sequence matched quiescent regions, and DNVs from healthy individuals from Genomes of The Netherland (GoNL) study overlapping fetal brain specific enhancers, are represented as blue dots. The size of the dot indicate observed number of DNVs. Black dotted line indicates *p*-value threshold of 0.05 while red dotted line indicate *p-*value after multiple test correction (*q-*value). b) Enrichment of context dependent tolerance score (CDTS) across fetal brain specific and human gain enhancers. X-axis indicates bins of CDTS score. Y-axis indicates enrichment of CDTS relative to whole genome control. The blue dots indicate human gained enhancers; green dots indicate fetal brain specific enhancers while orange dots indicate sequence matched control regions. b) Number of de novo variants (DNVs) constrained enhancer (turquoise) and sequence matched control regions (orange).

Despite that DNVs in ID (but not from healthy individuals) showed greatest enrichment in fetal brain enhancers compared to enhancers from other tissues or control regions, this result was not significant. We next explored whether subset of fetal brain enhancers that are under population constrain show enrichment for DNVs from ID patients.

Although variants in enhancer regions can lead to severe developmental defects^40^, functional redundancy of enhancers of developmental genes reduces the likelihood of severe functional consequences of enhancer variants^41^. Thus, disease-causing variants should specifically occur in enhancers that are intolerant to variants within human populations. The recently developed context dependent tolerance score (CDTS) provides sequence constraints across human population in non-coding regions of the genome at 10bp resolution^42^. The human gain enhancers as well as fetal brain-specific enhancers showed significant enrichment (*P* < 2.2×10^-16^ and *P* < 2.2×10^-16^ respectively) for constrained genomic regions (CDTS <=30; **Figure 1b)** when compared to sequence composition-matched control regions. This finding suggests that the putative fetal brain enhancers tend to be intolerant to variants within human populations. A total of 25 fetal brain enhancer DNVs were located within constrained regions (CDTS score <= 30) of the genome. We found marginal enrichment of DNVs in constrained enhancers (CDTS<=30) as compared to control quiescent regions (*P*=0.072; **Figure 1c**). However, when we restricted the analysis to highly constrained regions (CDTS<=20), we found significant excess of DNVs (*P*=0.027; **Figure 1c**) in fetal brain enhancers as compared to control regions. The increased burden of DNVs in ID patients in enhancers that are intolerant to variants within the human population suggests that DNVs in population-constrained regions are more likely to be functional.

### DNV-containing enhancers were associated with ID-relevant genes

We next investigated the hypothesis that DNVs are preferentially located in enhancers that are connected with genes that are plausible etiological mediators of ID. To identify potential target genes of fetal brain enhancers we used the following datasets in sequential order: promoter capture Hi-C (PCHi-C) data^31^ from neuronal progenitor cells (NPC); correlation of H3K27ac ChiP-seq signal at promoters and enhancers across multiple tissues; and promoter-enhancer correlation using chromHMM segmentation data. The closest fetal brain expressed gene was assigned as a target gene for 24% of the enhancers that remained unassigned after application of these approaches. For all approaches, we restricted our search space to brain topologically associated domains (TADs)^43^ as the majority of enhancer-promoter interactions happen within TADs^44^.

Next, we compiled genes that have previously been implicated in ID or related neurodevelopmental disorders, using three gene sets; known ID genes^4, 45^, genes implicated in neurodevelopmental disorders in the Deciphering Developmental Disorder (DDD) project^32^ and autism risk genes (SFARI genes)^46^. This provided us with a unique set of 1,868 genes previously implicated in neurodevelopmental disorders. The genes that were connected with DNV-containing fetal brain enhancers were enriched for known neurodevelopmental disorder genes (25 genes, *P* = 0.025; **Table S5**).

We further observed that the target genes of DNV-containing enhancers were not only involved in nervous system development (*P* = 7.4×10^-4^; **Table S6**) but also predominantly expressed in the prefrontal cortex (*P* =6.5 x10^-3^; **Table S7**), a brain region that has been implicated in social and cognitive behavior, personality expression, and decision-making.

The potential functional effect of heterozygous enhancer variants is expected to be mediated through altered expression of target genes. Recently, it has been shown that the majority of known severe haploinsufficient human disease genes are intolerant to loss of function (LoF) variants^47^. We compared the putative target genes of DNV-containing enhancers with the recently compiled list of genes that are intolerant to LoF variants (pLI >=0.9)^47^. We found that a significantly higher proportion of enhancer DNV target genes were intolerant to LoF variants than expected (*P* = 4.2×10^-5^; **Table S8**).

Taken together, our analysis shows that heterozygous DNVs are predominantly found in enhancers that are connected with genes that show preferential expression in the pre-frontal cortex, have been previously implicated in ID or related disorders, and exhibit intolerance to LoF variants.

### Recurrently mutated enhancer clusters

We did not identify individual enhancers that are recurrently mutated (contains two or more DNVs from unrelated probands), but investigated whether clusters of enhancers that regulate the same gene show recurrent DNVs. We defined the clusters of enhancers as the set of enhancers that are connected with the same gene. The enhancer clusters associated with three genes, *CSMD1*, *OLFM1* and *POU3F3* were recurrently mutated with two DNVs in each of the enhancer clusters **(Figure S1).** A presence of three enhancer clusters with recurrent DNVs within the cohort of 47 ID probands was significantly higher than expected (pervariant test *P* = 0.016). All three genes (*CSMD1*, *OLFM1* and *POU3F3)* play a role in nervous system development^48–50^. Heterozygous variants in *POU3F3* protein coding regions have been recently implicated in ID^51, 52^. A known role of these genes in nervous system development and the presence of recurrent variant in enhancer clusters associated with these genes in ID cohort suggest that these enhancer DNVs may contribute to ID.

### Functional disruption of enhancer function by ID DNVs

Enhancers regulate gene expression through the binding of sequence-specific transcription factors (TFs) at specific recognition sites ^53^. DNVs could elicit phenotypic changes because they alter the sequence of putative TF binding sites or because they create putative TF binding sites that have an impact on target gene expression. We used stringent criteria for TF motif prediction as well as motif disruption (see Methods). Of the 82 *de novo* SNVs that were located in fetal brain enhancers, 32 (39%) were predicted to alter putative TFBS affinity, either by destroying or creating TFBS **(Table S9a**). In comparison only 23.5% of the DNVs from healthy individuals (GoNL) located in fetal brain enhancers were predicted to alter putative TFBS affinity (**Table S9b**). Thus, a significantly higher proportion of the fetal brain enhancer DNVs from ID probands found to disturb putative TFBS affinity as compared to GoNL DNVs (Fisher’s exact test *p*-value = 0.0067; **Table S9c**). Furthermore, the fetal brain enhancer DNVs from ID probands frequently disturbed putative binding sites of TFs that were predominantly expressed in neuronal cells (*P* = 0.022; **Table S9d**), whereas no such enrichment was observed for GoNL DNVs (**Table S9e).** Taken together our results suggest that the enhancer DNVs from ID probands were more likely to affect the binding sites of neuronal transcription factors, and could influence the regulation of genes involved in nervous system development through this mechanism.

To test the functional impact of regulatory variants on enhancer activity, we selected 11 potential disease-causing enhancer DNVs **(Table S10),** and investigated their functional impact in luciferase reporter assays in the neuroblastoma cell line SH-SY5Y. Of the 11 enhancers containing DNVs, 10 showed significantly higher activity than the negative control in at least one allelic version, indicating that they do indeed function as active enhancers in this neuronal cell line (**Figure 2**). Amongst these 10 active enhancers, nine showed allele-specific activity, with five showing loss of activity and four showing gain of activity of the DNVs **(Figure 2)**. Eight out of nine DNVs that showed allele-specific enhancer activity altered a core base of the TF motif with position specific weight >= 0.95, and thereby disrupted the predicted affinity of the cognate TF **(Table S9)**. The *CSMD1* enhancer cluster had two DNVs (chr8: 2177122C>T and chr8: 2411360T>C) in two unrelated ID probands (Family 6 and Family 3 respectively). Both of these showed gain of activity compared to the wild type allele (**Figure 2**). By contrast, two DNVs in the OLFM1 enhancer cluster (chr9:137722838T>G and chr9:137333926C>T) from two unrelated ID probands (Family 4 and Family 12 respectively) caused loss of activity (**Figure 2**). These results demonstrate that selected DNVs from ID patients in fetal brain enhancers alter predicted TF binding affinity and have a functional impact on enhancer activity assays.

**Figure 2:**
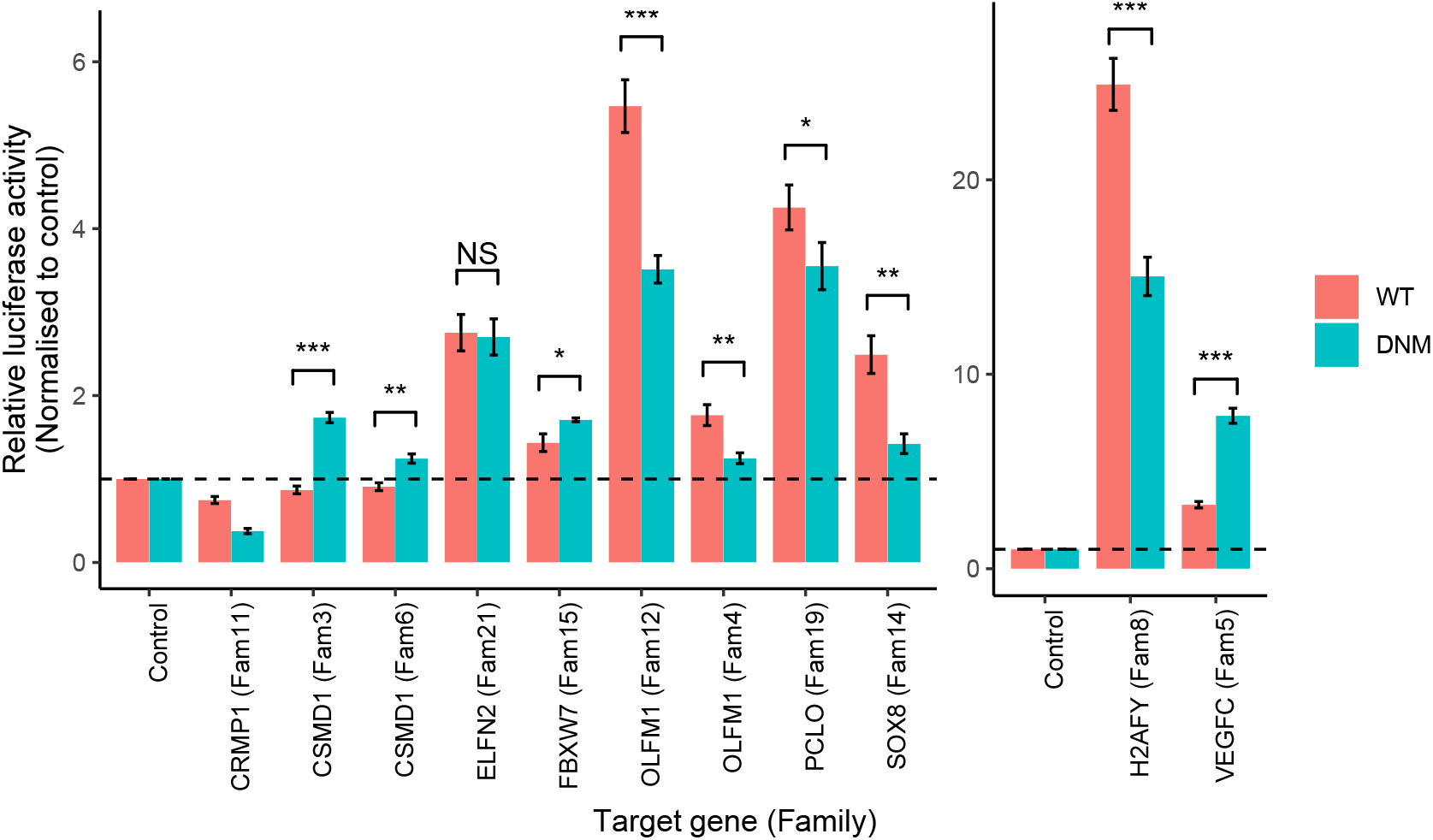
Effect of DNV on enhancer activity. Duel luciferase reporter assay of wild type (reference) and the mutant (DNV) allele. X-axis indicates putative target genes of the enhancer, while the family IDs are shown in brackets. Y-axis indicates relative luciferase activity normalised to empty plasmid. The error bars indicate standard error of means of three biological replicates. The enhancers associated with genes H2AFY and VEGFC are plotted with separately with different Y-axis scale because of high activity of these enhancers. The significance level was calculated using two-tailed t-test. *** Indicates p-value <=0.001, ** indicate p-value between 0.01 and 0,001 while * indicates p-value between 0.01 to 0.05.

### Function of the *CSMD1* enhancer in a neural differentiation model

We further explored the possible functional relevance of DNVs in the *CSMD1* enhancer cluster. Both ID probands with these variants showed developmental delay and both were overgrown with high birth weights (above the 91st centile) and remain large throughout postnatal life. The *CSMD1* gene is involved in neurogenesis^54^, and intronic variants have been shown to be associated with schizophrenia^55^. We focused on DNV chr8:2411360T>C from Family 3, because motif analysis predicted that it disrupts the binding site for the transcriptional repressor *TCF7L1*^56, 57^ (**Figure 3a**), which is known to inhibit premature neurogenesis^58^. The enhancer that harbors this DNV is located 2.4 Mb away from the *CSMD1* promoter (**Figure 3a**). To test the function of this enhancer, we performed lentiviral CRISPR interference (CRISPRi) by recruiting dCas9 fused with the repressor KRAB domain in the Lund human mesencephalic (LUHMES) neuronal precursor cell line, which do not express *CSMD1* mRNA **(Figure 3d),** and then differentiated LUHMES cells into neurons. We found that CRISPRi of the *CSMD1* enhancer led to significantly higher expression of *CSMD1* in neurons than in non-targeted control cells (*P*=0.004; **Figure 3b**). Given that the KRAB domain represses transcription through heterochromatin spreading^59, 60^ or simply by steric interference of endogenous regulatory components^61^, this result suggests that CRISPRi of this enhancer prevented binding by a transcriptional repressor, and thereby enabled overexpression of *CSMD1*.

**Figure 3:**
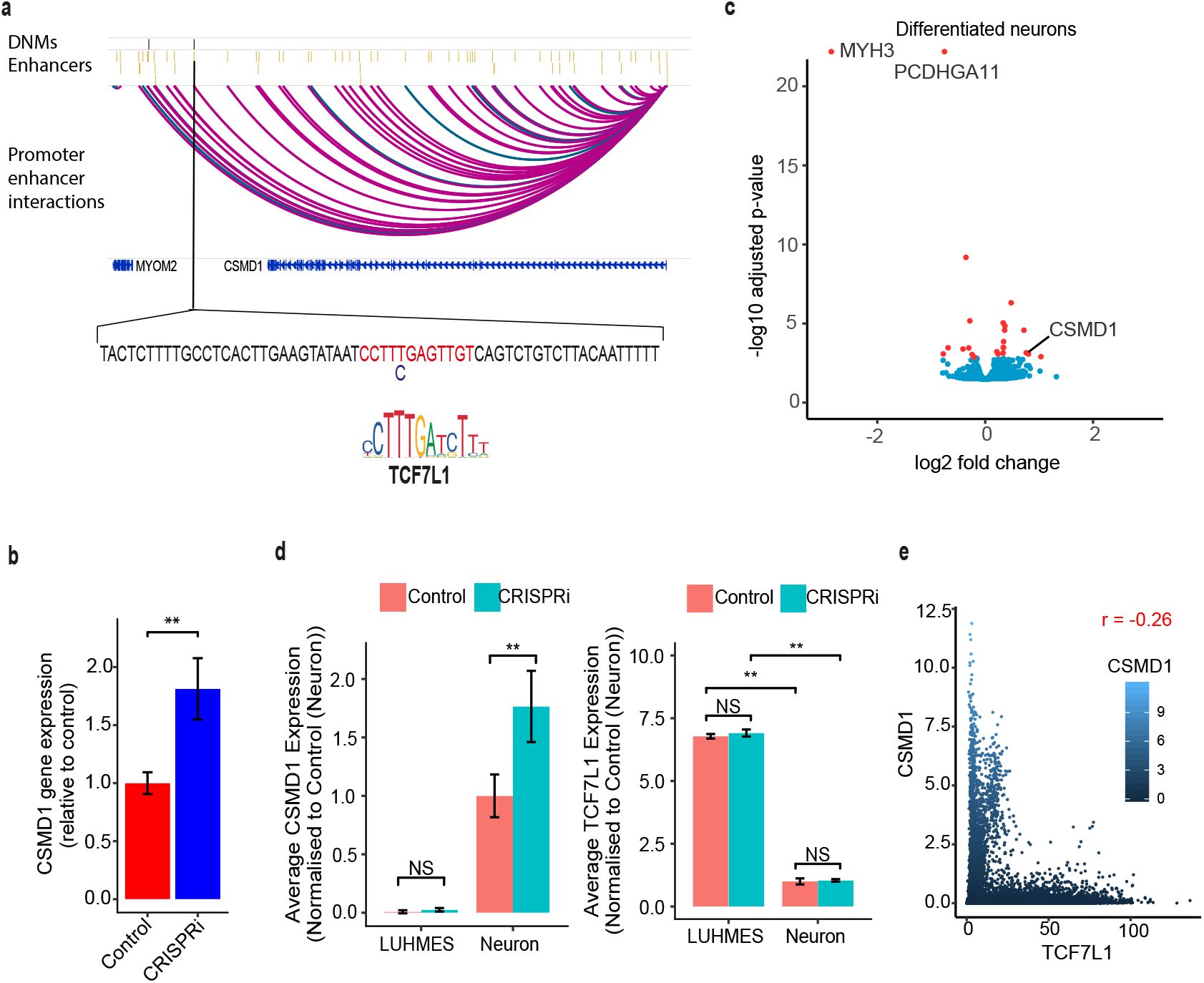
*Do novo* variant in CSMD1 enhancer lead premature activation of CSMD1. a) Enhancer-promoter interactions in *CSMD1* enhancer cluster. Pink arcs represent fetal brain specific enhancer *CSMD1* promoter interactions while green arcs represent human gain enhancer-promoter interactions. *De novo* variant in *CSMD1* enhancer (T>C) alters the core base of the *TCF7L1* transcription factor binding site. b) *CSMD1* expression measured using qPCR. CSMD1 shows significantly higher expression in cells containing KRAB-dCas9 (blue bar) as compared to control wild type cells (red bars). The error bars indicate the standard error of means of three replicates. The significance level was calculated using a two-tailed t-test. c) Volcano plot depicting differentially expressed genes between *CSMD1* enhancer CRISPRi and the control in differentiated neuron using RNA-seq. The red dots indicate significantly differentially expressed genes. d) Gene expression of *CSMD1* and *TCF7L1* genes in LUHMES cells and differentiated neurons, as determined by RNA-seq. e) Dot plot indicating gene expression pattern of *CSMD1* and *TCF7L1* across multiple tissues. Gene expression data was obtained from GTEx database.

Because the DNV disrupts a high-affinity binding sequence for TCF7L1, the CRISPRi experiments suggest that TCF7L1 could be a negative regulator of *CSMD1* during neuronal differentiation. Consistent with this model, *TCF7L1* mRNA was highly expressed in LUHMES cells, which do not express *CSMD1*, and was downregulated upon differentiation to neurons, which do express *CSMD1* **(Figure 3d)**. Further analysis of GTEx^62^ showed that the expression of *CSMD1* and *TCF7L1* are almost mutually exclusive across human tissues **(Figure 3e)**. Taken together, these results indicate that the candidate causal DNV disrupts a TCF7L1 recognition sequence in an enhancer that acts as a negative long range regulator of *CSMD1* during neural differentiation.

To investigate the impact of perturbing the *CSMD1* enhancer on neural differentiation, we performed RNAseq upon differentiation of neuronal precursors to neurons. RNAseq data confirmed significant up-regulation of CSMD1 in CRISPRi inactivated neurons as compared to controls. In addition, the genes *MYH3* (myosin heavy chain 3), expressed exclusively during embryonic development^63^, and *PCDHGA11* (Protocadherin Gamma-A11), which plays a significant role in establishment of cell-cell connections in the brain^64^.ac, showed a significant strong downregulation **(Figure 3c)**. These experiments indicate that the enhancer that harbors the regulatory chr8:2411360T>C variant modulates critical regulators of axon formation and neuronal connectivity.

### *CSMD1* enhancer cluster DNVs and ASD

The genetic underpinnings of both ID and autism spectrum disorders (ASD) have been ascribed to abnormal neuronal development, and may share molecular mechanisms^65, 66^. Approximately 10% of individuals with ID have ASD, while 70% of individuals with ASD have some level of ID^66^. Therefore, we explored the occurrence of DNVs in the *CSMD1* enhancer cluster in large-scale GS studies on ASD. The Simons Simplex Collection (SSC) has sequenced GS of 1,902 ASD families (quads)^11^, while the MSSNG database has compiled GS of 2,281 ASD trios^37^. Within the SSC cohort, four DNVs from four unrelated ASD cases were located in the *CSMD1* enhancer cluster, while no DNVs in the *CSMD1* enhancer cluster were found in 1,902 unaffected siblings. Additionally, we found four ASD patients from the MSSNG database harboring DNVs in the *CSMD1* enhancer cluster. Thus, we identified a total of eight DNVs in ASD cohorts. Recurrent variants in the *CSMD1* enhancer cluster in ID and ASD, but not in unaffected siblings, further reinforce the possible role of *CSMD1* enhancer variants in these neurodevelopmental disorders.

## Discussion

Despite the recent widespread use of genome sequencing, the true burden of disease-causing variants in enhancers is unknown. This is largely due to an inability to predict the pathogenicity of enhancer variants based on sequence features. Aggregation of a minority of disease-causing variants with the majority of benign regulatory variants nullifies any signal from disease-causing variants in non protein-coding genomic regions in disease cohorts. It is noteworthy that in protein coding regions of the genome only protein-truncating variants, but not other protein-coding variants, show significant enrichment in neurodevelopmental disorders^11, 67^. The analysis of DNVs in selected monogenic phenotypes provides a powerful analysis instrument, because it can focus on a relatively small number of variants that have increased likelihood of being disease-causing. In this study, we show that DNVs in a cohort of patients with ID exhibit a non-random genomic distribution that differs from DNVs observed in healthy individuals, with several features that are consistent with a disease-causing role of noncoding DNVs. DNVs from patients with ID were thus selectively enriched in fetal brain enhancers that exhibit variantal constraints in humans, in enhancers associated with genes that are ID-relevant, intolerant to loss of function variants and specifically expressed in pre-frontal cortex, and in disease-relevant transcription factor binding sites. Our studies further provide experimental evidence indicating that ID DNVs are enriched for regulatory variants.

Nearly half of all human enhancers have evolved recently^68^, and advanced human cognition has been attributed to recently evolved human specific brain enhancers^12^. We show that fetal brain enhancers show selective constraint in humans **(Figure 1b),** and also found that DNVs in patients with ID are significantly enriched in fetal brain enhancers that are constrained in human populations. Enrichment of DNVs in enhancers that are intolerant to variants within human populations suggests that the DNVs in such essential enhancers are more likely to be functional. Consistent with this prediction, we show that DNVs from ID patients disturb binding sites of neuronal transcription factors within fetal brain enhancers more frequently than DNVs from healthy individuals. Furthermore, more than 90% of the DNVs tested showed allele-specific activity (**Figure 2).** Genetic and experimental observations, therefore, indicate that DNVs from patients with ID are enriched in functional neural regulatory variants.

The identification of genes that are recurrently mutated across multiple disease individuals has been a major route to discover novel disease genes^69^. We identified recurrent variants within three fetal brain enhancer clusters associated with genes involved in nervous system development (*CSMD1, OLFM1, POU3F3*), and found that this enrichment was significant relative to expectations. All three genes show high pLI score (1, 1, 0.88 respectively), indicating that they are intolerant to loss of function variants and dosage sensitive, and one of them (*POU3F3)* was recently shown to harbor disease-causing heterozygous variants in patients with ID^52^. Our results suggest that enhancer variants could lead to dysregulation of these genes during nervous system development and thereby contribute to the etiology of ID.

Among these three loci, we focused on *CSMD1*, a gene that is highly expressed in the central nervous system, particularly in the nerve growth cone^50^. Common genomic variants in *CSMD1* are associated with schizophrenia and neuropsychological measures of general cognitive ability and memory function^70–72^. Furthermore, *CSMD1* knockout mice show strong neuropsychological defects^73^. Our motif sequence analysis, luciferase assays, and CRISPRi experiments indicate that an ID DNV in this locus affects an enhancer that acts as a negative regulator of *CSMD1,* and impacts expression of other neuronal genes. We further identified enrichment of variants in the *CSMD1* enhancer cluster in cohorts of patients with neurodevelopmental disorders that have been proposed to share pathogenic mechanisms. Collectively, these findings implicate *CSMD1* as a strong candidate target gene for ID disease-causing regulatory variants.

Taken together, our work has integrated whole genome sequences, epigenomics and functional analysis to examine the role of regulatory DNVs in ID. Despite the genetic heterogeneity of ID, which severely hampers efforts to unequivocal demonstrate a causal role for individual noncoding variants, our results provides multiple lines of evidence to indicate that functional regulatory variants in selectively constrained, stage-specific brain enhancers contribute to the etiology of ID. This work should prompt extensive genetic analyses and variant-specific experimental modeling to elucidate the precise role of regulatory variants in ID.

## Acknowledgments

We thank the families of the affected children for their time and support for the research. This research was supported by the National Institute for Health Research (NIHR) Imperial Biomedical Research Centre. This work was funded by grants from the Wellcome Trust Institute Strategic Support and National Institute for Health Research (NIHR) Imperial Biomedical Research Centre, Institute for Translational Medicine and Therapeutics (P70888) obtained by S.S.A.. J.F.’s work was funded by grants from the Wellcome Trust (WT101033 to J.F.), Medical Research Council (MR/L02036X/1 to J.F.), European Research Council Advanced Grant (789055 to J.F.). M.M.P.’s work was funded by the Wellcome Trust Research seed award and funding from the School of Medicine and Dentistry, QMUL. T.N.K. was partially supported by Government of Pakistan under PSDP project “Development of National University of Medical Sciences (NUMS), Rawalpindi”. We thank Prof Andrew Jackson for helpful discussions and obtaining ethical approval for the study. We thank Mrs Sophie Shi for contributing to reagent generation. We also thank Dr Patrick Short and Dr Kaitlin Samocha both from Sanger Institute for providing trinucleotide probability table and helpful discussion on mutational model, respectively.

## Author contributions

Conceptualization, S.S.A.; Acquiring funding, S.S.A.; Patient recruitment and sample collection, A.K.L, W.W.L. and D.R.F.; Bioinformatics analysis-Variant calling, S.S.J. and D.P.; Bioinformatics analysis: Enhancer-promoter interactions, G.A. and S.S.A.; Bioinformatics analysis: All other: S.S.A.; Experiments – Sanger sequencing: D.M.; Experiments – Luciferase: M.G.G., T.K., M.G.D., I.C., S.Z. and W.C.; Experiments – CRISPR: M.G.D.; Writing – Original Draft: S.S.A.; Writing – Review & Editing, J. F., M.M.P., M.G.D., I.C. and M.G.G.; Supervision, D.R.F., J.F., M.M.P. and S.S.A.

## Data availability

The whole genome sequence data as well as variant calls that support the findings of this study are available on request from the corresponding author [S.S.A.]. The data are not publicly available because it contains information that could compromise research participant privacy/consent. The RAN-seq data has been deposited to ArrayExpress with accession number E-MTAB-8027. RNA-seq data can be accessed using following URL https://www.ebi.ac.uk/arrayexpress/experiments/E-MTAB-8027/

---

**Figure S1:**
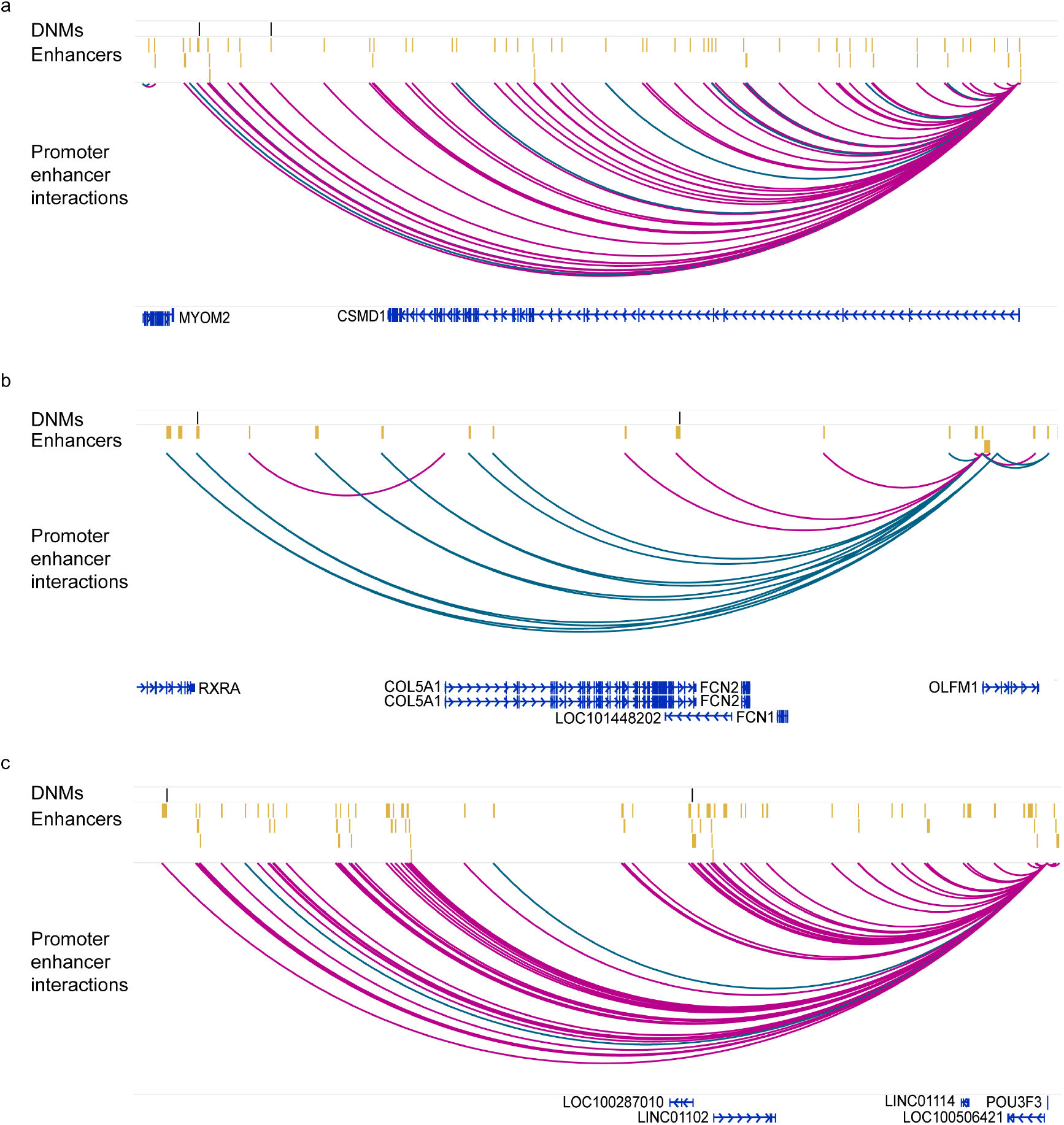
Recurrent de novo mutations (DNMs) in enhancer clusters: a) Recurrent DNMs in CSMD1 enhancer cluster. b) Recurrent DNMs in OLFM1 enhancer cluster. c) Recurrent DNMs in POU3F3 enhancer cluster. Black lines indicate DNMs while yellow bars indicate enhancers. Pink arcs represent fetal brain specific enhancer-promoter interactions while green arcs represent human gain enhancer-promoter interactions.

